# Progastrin activates colonic fibroblasts and induces a paracrine pro-migratory program in colorectal cancer cells

**DOI:** 10.64898/2026.04.12.718027

**Authors:** Nicolas Fénié, Timothy C. Wang, Serge Roche, Audrey Ferrand

**Affiliations:** INSERM UMR1037, Cancer Research Center of Toulouse (CRCT), Université Paul Sabatier Toulouse III, Toulouse, France; Division of Digestive and Liver Diseases, Irving Cancer Research Center, Columbia University, New York, NY, USA; Equipe Labellisée LIGUE 2020 and FRM 2023, CRBM, Univ Montpellier, CNRS, Montpellier, 34293, France; University of Toulouse, INSERM, I2MC, Toulouse, France

**Keywords:** progastrin, colorectal cancer, cancer-associated fibroblasts, tumour-stroma crosstalk, migration, CXCL12, CXCL8

## Abstract

**Purpose:** Progastrin, aberrantly expressed in colorectal cancer (CRC), is an established trophic factor for tumour epithelial cells. Whether it also promotes CRC progression by reprogramming stromal fibroblasts remains unclear. We investigated progastrin-induced colonic fibroblast activation and its functional consequences on CRC cell migration.

**Methods:** Fibroblast activation was assessed in the colonic mucosa of hGAS mice and in the human normal colonic fibroblast line CCD18Co exposed to synthetic progastrin. The impact of tumour-derived progastrin on epithelial cell motility was analysed using HCT116 cells expressing a control shRNA (shLuc) or a progastrin-targeting shRNA (shPG) in transwell migration assays performed with or without fibroblasts. Candidate paracrine mediators were evaluated by RT-qPCR, ELISA and neutralization experiments, and signalling was interrogated using the PI3K inhibitor LY294002.

**Results:** Colonic fibroblasts from hGAS mice displayed stromal FAPα and αSMA expression, indicating fibroblast activation *in vivo*. In CCD18Co cells, progastrin increased FAPα and αSMA protein levels. Fibroblasts enhanced HCT116 cell migration. This effect was stronger when tumour cells expressed progastrin or when fibroblasts were preconditioned by progastrin-producing HCT116 cells. Progastrin induced CXCL12/SDF-1 and CXCL8/IL-8 expression and secretion by fibroblasts, and neutralization of either chemokine abrogated the additional migratory effect conferred by progastrin-activated fibroblasts. Progastrin triggered sustained Akt phosphorylation in fibroblasts, while PI3K inhibition suppressed CXCL12 and CXCL8 secretion and abolished fibroblast-dependent tumour cell migration.

**Conclusion:** These data identify a stromal dimension of progastrin signalling in CRC and support a model in which tumour-derived progastrin activates colonic fibroblasts and elicits a PI3K/Akt-dependent paracrine programme that enhances CRC cell migration.

## Introduction

Colorectal cancer (CRC) remains a major global health burden. According to GLOBOCAN 2022 estimates, it is among the most frequently diagnosed malignancies worldwide and the second leading cause of cancer-related death [1]. Although localized tumours can be cured by surgery, the prognosis changes profoundly once tumour cells acquire the capacity to invade adjacent tissues and disseminate to distant organs. Metastatic CRC therefore remains the clinically decisive stage of the disease and continues to account for most CRC-related deaths [2,3].

The biological transition from local tumour growth to invasive and metastatic competence is not governed by epithelial alterations alone. It is increasingly understood as a microenvironment-dependent process in which stromal cells actively shape tumour behaviour. Among stromal populations, cancer-associated fibroblasts (CAFs) are now recognized as major regulators of extracellular matrix remodelling, paracrine signalling, tumour cell plasticity and disease progression. In CRC, fibroblast-rich tumours, particularly those assigned to the mesenchymal CMS4 subtype, are associated with poor outcome [4]. Experimental studies further indicate that CRC cells activate adjacent fibroblasts, which in turn enhance tumour cell migration and invasion [5], while stromal TGF-β programmes promote metastasis initiation [6]. More recently, patient-derived CAFs have been shown to drive cancer cell invasion already at the earliest stages of colorectal tumorigenesis [7].

Within this framework, progastrin is a compelling candidate mediator of tumour-stroma crosstalk. The GAST gene is a downstream target of the Wnt/β-catenin/TCF-4 pathway [8], and oncogenic Ras induces gastrin gene expression in colon cancer cells [9]. Cooperation between Ras and β-catenin further enhances gastrin promoter activity [10]. As a consequence of these early oncogenic alterations, CRC epithelial cells frequently express gastrin gene; however, because they incompletely process the prohormone, they predominantly produce progastrin rather than amidated gastrin [11,12]. Circulating progastrin is increased in patients with CRC, further supporting its pathological relevance [13].

The tumour-promoting role of progastrin in CRC has been documented mainly from the perspective of epithelial biology. Gastrin gene expression is required for the proliferation and tumorigenicity of colon cancer cells [14]. In transgenic mice overexpressing human progastrin, the colonic mucosa displays hyperproliferation and increased susceptibility to carcinogen-induced neoplasia [15–17]. In tumour cells, progastrin contributes to maintenance of stem-like properties and is required for the tumorigenic potential of CD133-positive CRC cells [18,19]. Progastrin also perturbs epithelial junctional organization [20], and blockade of extracellular progastrin reduces the migratory and invasive properties of CRC cells [21]. These observations suggest that progastrin may contribute not only to tumour growth, but also to invasion-related phenotypes. Nevertheless, its stromal effects remain incompletely defined.

A first indication that progastrin can act on mesenchymal cells came from Duckworth et al., who showed that progastrin stimulates colonic myofibroblasts to secrete IGF-II and thereby indirectly enhances epithelial proliferation [22]. We reasoned that progastrin might more broadly reprogramme resident colonic fibroblasts toward a CAF-like, pro-migratory state. In the present study, we therefore investigated whether progastrin activates normal colonic fibroblasts, whether tumour-derived progastrin enhances fibroblast-dependent CRC cell migration, and which paracrine mediators and intracellular signalling pathways underpin this effect. We identify a PI3K/Akt-dependent fibroblast response to progastrin that drives CXCL12 and CXCL8 secretion and functionally promotes CRC cell migration.

## Materials and methods

### Animals

hGAS mice in an FVB/N background and corresponding control FVB/N littermates have been described previously [15]. Animals were maintained under standard conditions with a 12h light/12 h dark cycle, and experiments were performed during daytime. hGAS and control FVB/N mice were sex matched and colon tissues from at least four hGAS mice and four age-matched control littermates (22–24 weeks) were analysed. All animal procedures were approved by the local institutional animal care committee.

### Immunohistochemistry

Formalin-fixed, paraffin-embedded colon tissues were subjected to heat-induced epitope retrieval in citrate buffer. Sections were incubated overnight with primary antibodies. FAPα and αSMA were detected using the Polyview kit (Vector Laboratories), whereas CXCL12 and CXCL8 were detected with the Dako EnVision+ HRP system. Primary antibodies were: αSMA (Abcam, ab21027 for tissue staining; ab7817 for western blotting), FAPα (Abcam, ab53066), CXCL8 (Sigma-Aldrich, WH0003576M5), and CXCL12 (R&D Systems, MAB350).

### Cell culture and reagents

The human CRC cell line HCT116 and the human normal colonic fibroblast line CCD18Co were cultured at 37°C in a humidified atmosphere containing 5% CO2. HCT116 cells were maintained in DMEM (4.5 g/L glucose) supplemented with 10% fetal calf serum, whereas CCD18Co cells were cultured in DMEM/F12 supplemented with 10% fetal calf serum. Cells were serum-starved for 24 h before stimulation or coculture. Where indicated, CCD18Co fibroblasts were treated with 10^−8^ M of synthetic human progastrin ([23], provided by A. Shulkes, University of Melboune) and/or pretreated for 1 h with the PI3K inhibitor LY294002 (10 μM; Calbiochem).

### Protein extraction and western blotting

Cells were washed with PBS and buffer A (50 mM HEPES, pH 7.5, 150 mM NaCl, 10 mM EDTA, 10 mM Na4P2O7, 100 mM NaF, 2 mM Na3VO4) and lysed for 15 min at 4°C in buffer A containing 1% Nonidet P40, 0.5 mM PMSF, 20 μM leupeptin and 10 μg/mL aprotinin. Protein concentrations were determined by BCA assay. Equal amounts of protein were resolved by SDS-PAGE, transferred to PVDF membranes, blocked in milk or BSA according to antibody provider recommendations, and incubated overnight at 4°C with primary antibodies. Immunoreactive bands were detected using HRP-conjugated secondary antibodies and SuperSignal West Pico chemiluminescent substrate (Thermo Scientific). For signalling analyses, membranes were probed with antibodies recognizing total Akt and phospho-Akt (Ser473). GAPDH served as loading control.

### Generation of HCT116-shPG and HCT116-shLuc cells

Short hairpin RNA (shRNA) sequences targeting progastrin/GAST (GAAGAAGCCTATGGATGGA) or luciferase (shLuc) were cloned into the pSIREN-RetroQ vector. Retroviral transduction of HCT116 cells was performed as described previously [23]. Stable polyclonal populations were selected with puromycin (1 μg/mL). GAST knockdown was verified by RT-qPCR (Supplementary Fig. S1).

### Preparation of conditioned media

For tumour cell-conditioned media, HCT116-shLuc or HCT116-shPG cells were plated at 5 × 10^5^ cells/mL and cultured for 24 h before replacement with serum-free medium. After 20 h, supernatants were collected, centrifuged at 1000 rpm for 5 min, and mixed 1:1 with serum-free DMEM/F12 before incubation with CCD18Co fibroblasts. For fibroblast-conditioned media, CCD18Co cells were plated at 1 × 10^5^ cells/mL, grown for 24 h, then switched to serum-free medium for 20 h. Supernatants were collected, centrifuged at 1000 rpm for 5 min, and mixed 1:1 with serum-free DMEM before addition to HCT116 cells.

### RNA extraction and RT-qPCR

Total RNA was extracted using the RNeasy kit (Qiagen) and treated with DNase (Invitrogen). First-strand cDNA synthesis was performed from 1 μg RNA using the SuperScript system (Invitrogen). Quantitative PCR was performed with SYBR Green chemistry on an ABI StepOne Plus instrument. Relative expression was calculated using the 2^−ΔΔCt^ method with ACTB as reference gene. Primer sequences were as follows: CXCL12 forward 5′-TGCCAGAGCCAACGTCAA-3′ and reverse 5′-GCTACAATCTGAAGGGCACAGTT-3′; CXCL8 forward 5′-CTGGCCGTGGCTCTCTTG-3′ and reverse 5′-CTTGGCAAAACTGCACCTTCA-3′; ACTB forward 5′-GCGCGGCTACAGCTTCA-3′ and reverse 5′-CTTAATGTCACGCACGATTTCC-3′; GAS forward 5′-TCCATCCATCCATAGGCTTC-3′ and reverse 5′-CCACACCTCGTGGCAGAC-3′.

### ELISA

For chemokine quantification, cells were plated in 100mm dishes. After treatment, supernatants were collected, concentrated using Centricon devices (Millipore), and analysed using ELISA kits for CXCL12 and CXCL8 (R&D Systems) according to the manufacturer’s instructions. Chemokine concentrations were normalized to cell number.

### Transwell migration assays

CCD18Co fibroblasts were plated in the lower compartment of 12-well plates at 5 × 10^5^ cells/mL and serum-starved for 24 h. HCT116 cells were seeded in 8.0μm pore transwell inserts at 5 × 10^6^ cells per insert and allowed to adhere for 4h before the inserts were transferred to the fibroblast-containing wells. After an additional 4h, epithelial cells were switched to serum-free medium and allowed to migrate for 20h. Membranes were fixed with 4% paraformaldehyde, stained with crystal violet, imaged on a Nikon E400 microscope coupled to a Sony DXC950 camera, and quantified using Morpho Expert software. Where indicated, fibroblasts were preconditioned for 20h with HCT116-shLuc or HCT116-shPG cells before replacement of inserts and measurement of HCT116-shPG migration. For neutralization experiments, antibodies against CXCL12 or CXCL8 were added to fibroblast cultures during the migration assay.

### Statistical analysis

Data are presented as mean ± SEM. Statistical analyses were performed using GraphPad Prism. Pairwise comparisons were analysed using two-tailed Student’s t-tests. P values < 0.05 were considered statistically significant.

### Reporting support

ChatGPT (OpenAI) was used solely for grammatical and stylistic editing of manuscript text; it was not used for data analysis, figure generation, or interpretation of results.

## Results

### Progastrin induces a CAF-like activation phenotype in colonic fibroblasts

To determine whether progastrin can activate resident colonic fibroblasts *in vivo*, we first examined the colonic mucosa of hGAS mice, which overexpress human progastrin [15]. Fibroblast activation protein-α (FAPα) is a stromal serine protease broadly associated with activated fibroblasts in epithelial cancers [25], whereas αSMA is a hallmark of myofibroblastic differentiation and CAF-like activation [26].

Immunohistochemistry revealed stromal FAPα expression in hGAS colonic mucosa, whereas no staining was observed in control FVB/N tissue (Fig. 1a). Likewise, αSMA was readily detected in the stromal compartment of hGAS mice but not in control animals (Fig. 1a), indicating that chronic exposure to progastrin is associated with fibroblast activation *in vivo*.

**Figure 1.**
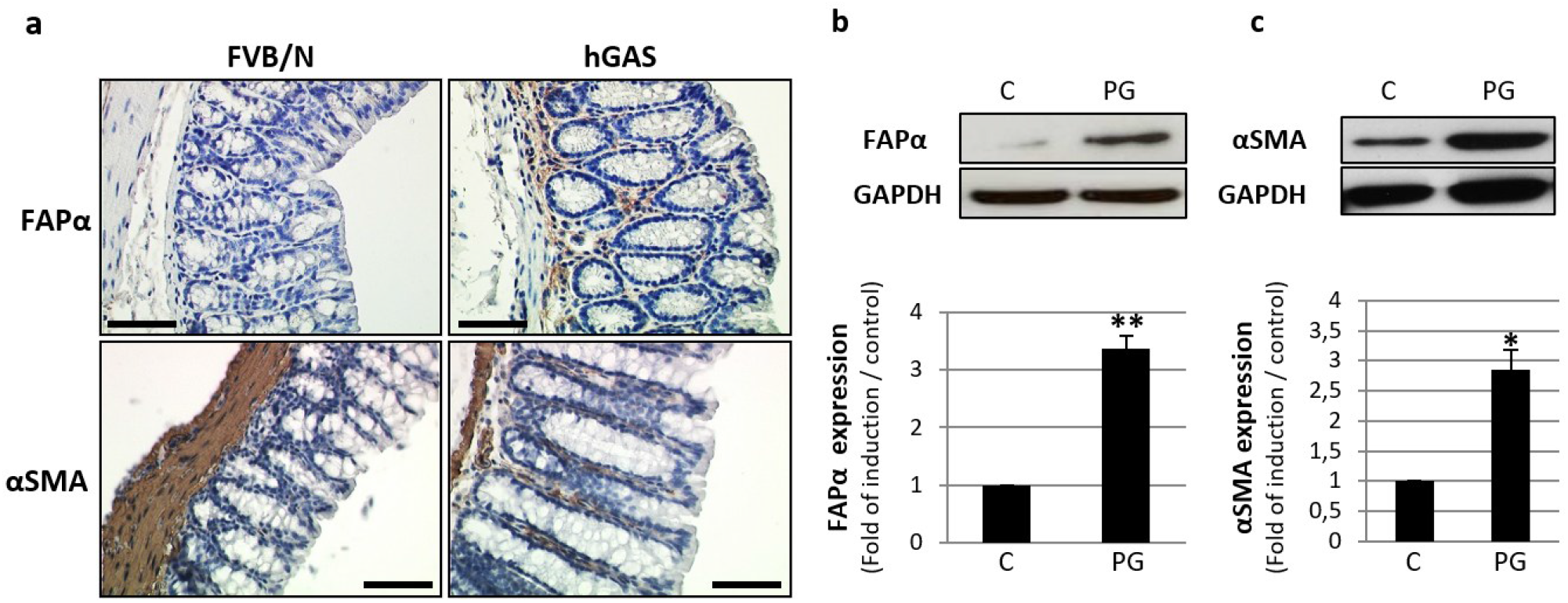
Progastrin induces a CAF-like activation phenotype in colonic fibroblasts. **a**, Representative immunohistochemistry for FAPα and αSMA in colonic mucosa from control FVB/N and hGAS mice (n=4; scale bars= 50µm). **b-c**, Western blot analyses of FAPα (n=4) and αSMA (n=3) expression in CCD18Co cells cultured in the absence (C) or presence (PG) of synthetic progastrin for 20 h, with corresponding densitometric quantification.

We next tested whether progastrin directly activates fibroblasts *in vitro*. In CCD18Co cells, 20h stimulation with synthetic progastrin increased FAPα and αSMA protein expression, as assessed by western blotting (Fig. 1b-c). Densitometric analysis showed a 3.10 ± 0.28-fold increase in FAPα (P = 0.0088; n = 4) and a 2.84 ± 0.34-fold increase in αSMA (P = 0.022; n = 3).

Together, these data indicate that progastrin is sufficient to drive a CAF-like activation programme in normal colonic fibroblasts.

### Tumour-derived progastrin enhances fibroblast-dependent HCT116 cell migration

We then asked whether fibroblast activation by tumour-derived progastrin has functional consequences for CRC cell motility. HCT116 cells, which endogenously produce progastrin [12], were engineered to express either a control shRNA (shLuc) or a progastrin-targeting shRNA (shPG). GAST knockdown reduced progastrin mRNA expression by 60 ± 1.2% (P = 0.00035; n = 3; Supplementary Fig. S1). We performed transwell migration assays in which HCT116 cells were seeded in the upper chamber, with or without CCD18Co fibroblasts plated in the lower well.

In transwell assays performed without fibroblasts, migration of HCT116-shLuc and HCT116-shPG cells did not differ, indicating that under these conditions the reduced progastrin expression did not measurably alter basal migration (Fig. 2). By contrast, the presence of CCD18Co fibroblasts strongly increased tumour cell migration. Migration increased by 3.34 ± 0.35-fold for HCT116-shPG cells (P = 0.0037; n = 5) and by 4.69 ± 0.26-fold for HCT116-shLuc cells (P = 0.00012; n = 5), with significantly greater migration of progastrin-expressing HCT116-shLuc cells than HCT116-shPG cells in the presence of fibroblasts (P = 0.001; Fig. 2).

**Figure 2.**
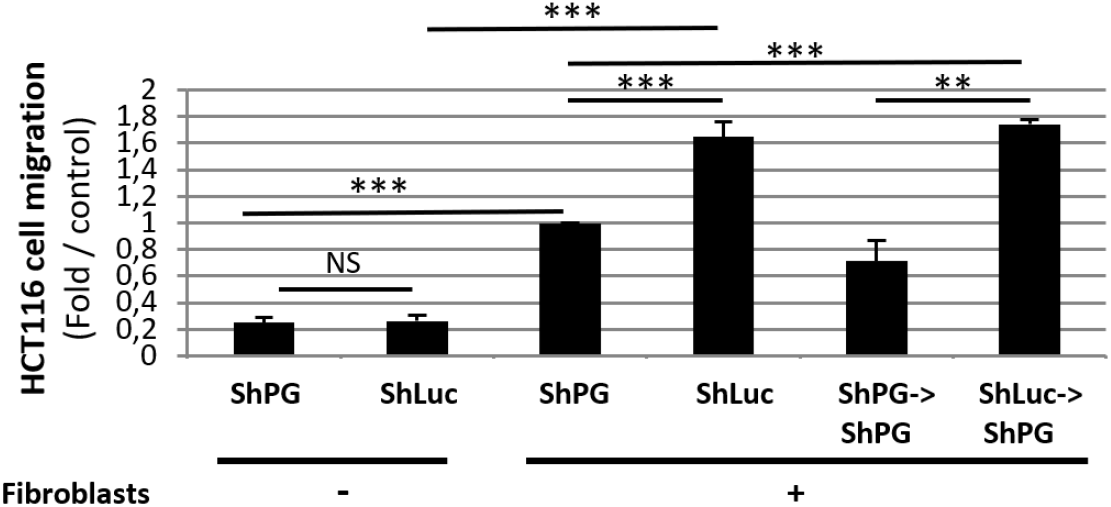
Tumour-derived progastrin enhances fibroblast-dependent migration of HCT116 cells. Migration of HCT116-shLuc and HCT116-shPG cells was analysed in transwell assays in the absence or presence of CCD18Co fibroblasts (n=5). In the preconditioning experiment, fibroblasts were first exposed to HCT116-shLuc or HCT116-shPG cells for 20 h, after which inserts were replaced and migration of HCT116-shPG cells was measured (n=3). Data are expressed as fold change relative to the indicated control conditions.

To test whether progastrin acts through fibroblast conditioning, CCD18Co cells were first exposed for 20 h to HCT116-shLuc or HCT116-shPG cells, after which inserts were replaced and migration of HCT116-shPG cells was measured. Fibroblasts preconditioned by HCT116-shLuc cells conferred a 2.66 ± 0.43-fold increase in HCT116-shPG migration compared with fibroblasts preconditioned by HCT116-shPG cells (P = 0.0079; n = 3; Fig. 2).

These results indicate that tumour-derived progastrin enhances CRC cell migration indirectly by overactivating stromal fibroblasts.

### Progastrin induces CXCL12/CXCL12 and CXCL8/CXCL8 expression and secretion by colonic fibroblasts

To identify mediators of the pro-migratory fibroblast response, we examined two chemokines with established roles in CRC progression: CXCL12/SDF-1 and CXCL8/IL8 [27,28].

When CCD18Co fibroblasts were cultured in conditioned medium from HCT116-shLuc cells, CXCL12 mRNA levels were 1.91 ± 0.21-fold higher than in fibroblasts exposed to conditioned medium from HCT116-shPG cells (P = 0.0029; n = 3; Fig. 3a). Direct stimulation of CCD18Co cells with progastrin also increased CXCL12 mRNA expression (2.25 ± 0.34-fold; P = 0.040; n = 3; Fig. 3b). Consistent with this transcriptional induction, CXCL12 secretion into fibroblast-conditioned medium increased 2.09 ± 0.19-fold after 20 h progastrin treatment (P = 0.032; n = 3; Fig. 3c). Increased stromal CXCL12 staining was also detected in hGAS mouse colonic mucosa compared with control tissue (Fig. 3d).

**Figure 3.**
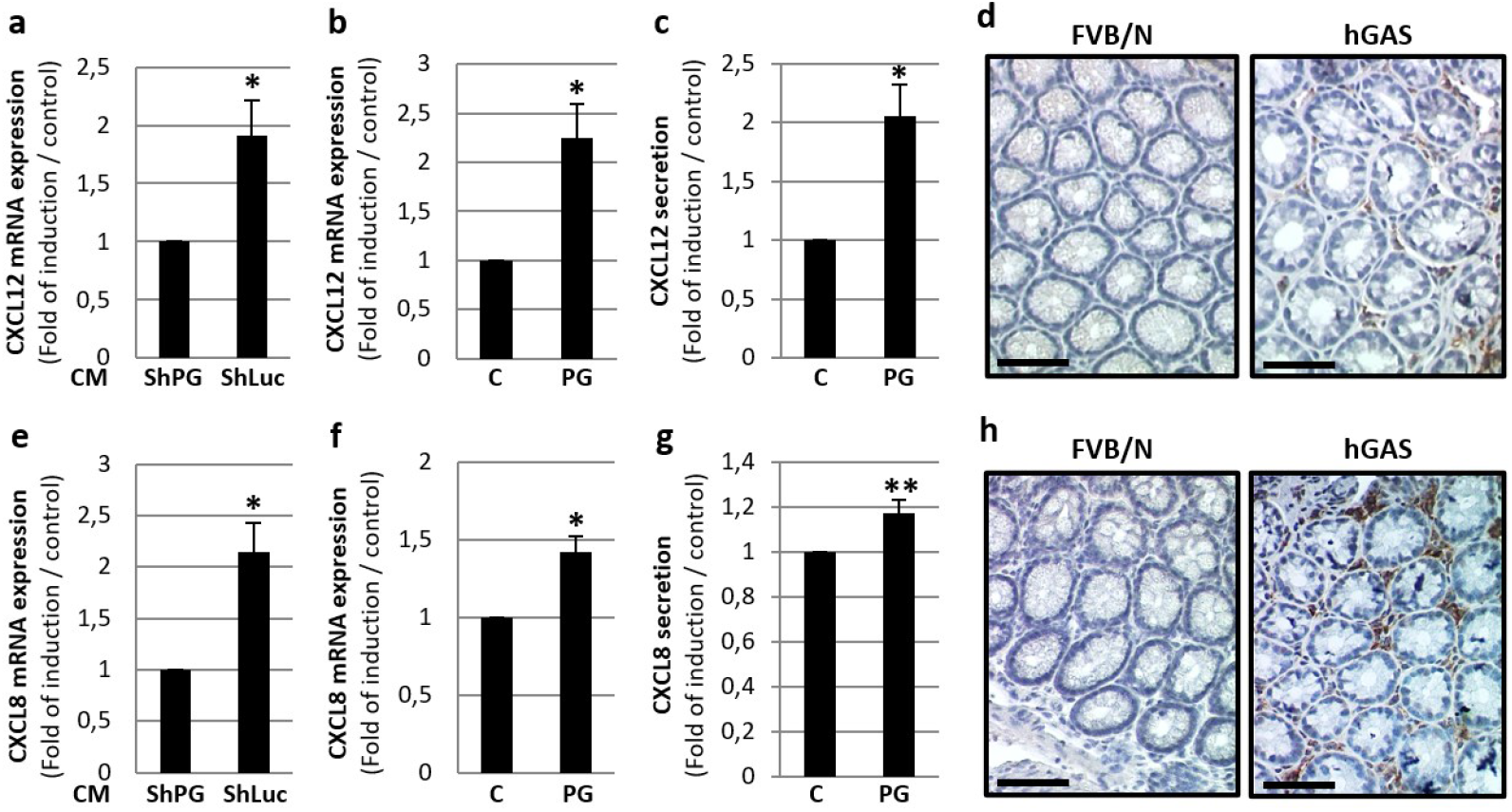
Progastrin induces CXCL12/SDF-1 and CXCL8/IL8 expression and secretion in colonic fibroblasts. **a**, RT-qPCR analysis of CXCL12 mRNA in CCD18Co fibroblasts cultured in conditioned medium from HCT116-shLuc or HCT116-shPG cells. **b**, RT-qPCR analysis of CXCL12 mRNA in CCD18Co cells stimulated directly with synthetic progastrin. **c**, ELISA quantification of CXCL12 secretion in conditioned medium from CCD18Co cells cultured in the absence (C) or presence (PG) of progastrin. **d**, Representative immunohistochemistry for CXCL12 in colonic mucosa from control FVB/N and hGAS mice (original magnification ×400). **e**, RT-qPCR analysis of CXCL8 mRNA in CCD18Co fibroblasts cultured in conditioned medium from HCT116-shLuc or HCT116-shPG cells. **f**, RT-qPCR analysis of CXCL8 mRNA in CCD18Co cells stimulated directly with synthetic progastrin. **g**, ELISA quantification of CXCL8 secretion in conditioned medium from CCD18Co cells cultured in the absence (C) or presence (PG) of progastrin. **h**, Representative immunohistochemistry for CXCL8 in colonic mucosa from control FVB/N and hGAS mice (scale bars= 50µm).

A similar pattern was observed for CXCL8. Compared with conditioned medium from HCT116-shPG cells, conditioned medium from HCT116-shLuc cells increased CXCL8 mRNA expression in CCD18Co fibroblasts by 2.14 ± 0.20-fold (P = 0.017; n = 3; Fig. 3e). Direct progastrin stimulation increased CXCL8 mRNA expression by 1.42 ± 0.09-fold (P = 0.040; n = 3; Fig. 3f) and CXCL8 secretion by 1.18 ± 0.03-fold (P = 0.0043; n = 4; Fig. 3g). *In vivo*, CXCL8 immunoreactivity was also increased in hGAS colonic mucosa (Fig. 3h).

These findings indicate that progastrin induces a fibroblast secretory programme enriched in CXCL12 and CXCL8.

### CXCL12 and CXCL8 mediate the additional migratory effect conferred by progastrin-activated fibroblasts

We next tested whether CXCL12 and CXCL8 are functionally required for the enhanced migration induced by progastrin-activated fibroblasts. Neutralizing antibodies against CXCL12 or CXCL8 were added to fibroblast cultures during transwell assays.

In HCT116-shPG cells, neither antibody significantly altered migration relative to the corresponding fibroblast-containing control condition (Fig. 4a). By contrast, both antibodies abolished the additional migratory advantage conferred by progastrin-activated fibroblasts to HCT116-shLuc cells, reducing migration to the level observed with HCT116-shPG cells (P = 0.047 for anti-CXCL12 and P = 0.022 for anti-CXCL8; Fig. 4a).

**Figure 4.**
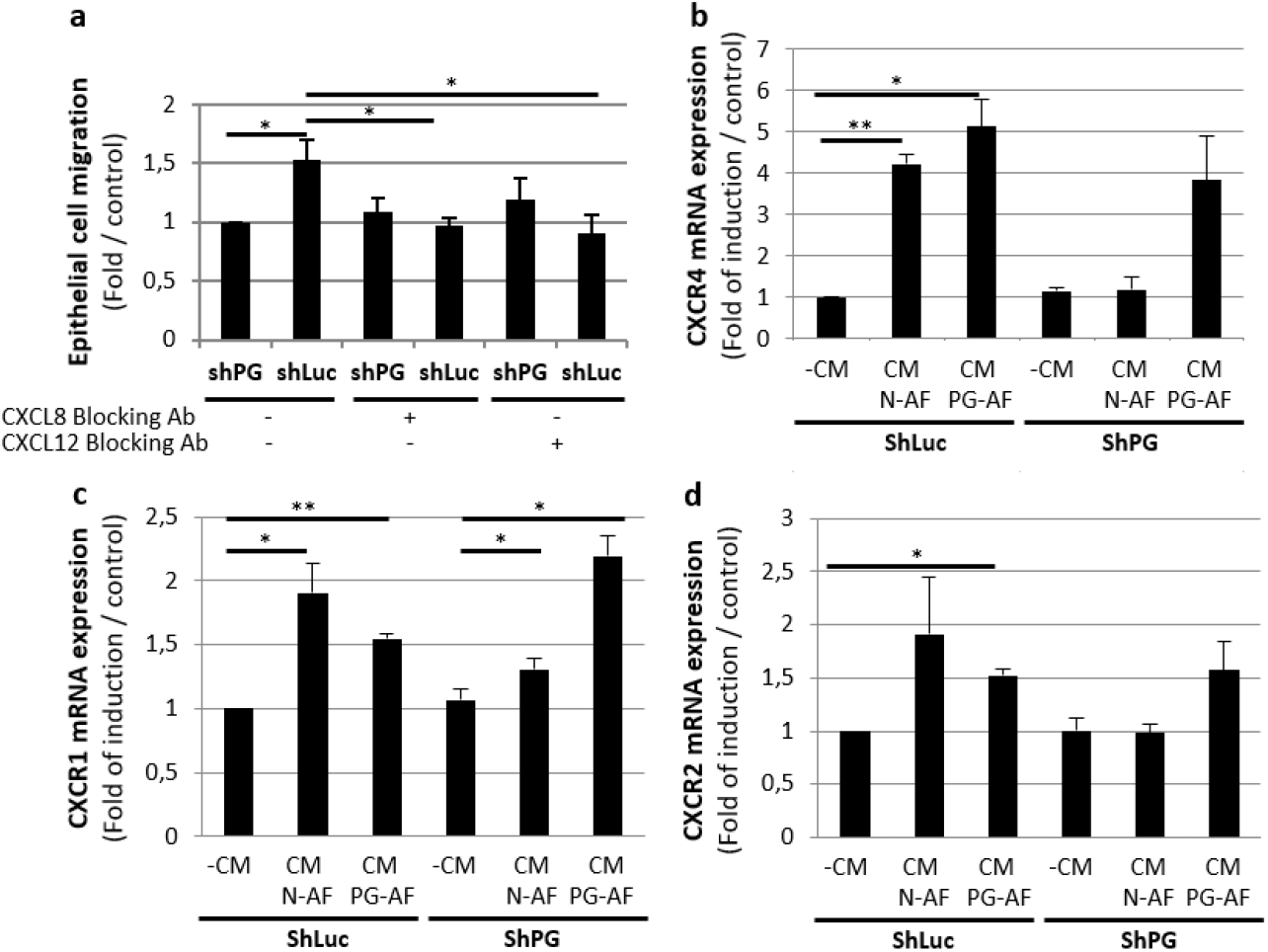
CXCL12 and CXCL8 mediate the pro-migratory effect of progastrin-activated fibroblasts. **a**, Transwell migration of HCT116-shLuc and HCT116-shPG cells in the presence of CCD18Co fibroblasts treated with neutralizing antibodies against CXCL12 or CXCL8. **b-d**, RT-qPCR analysis of CXCR4, CXCR1 and CXCR2 mRNA expression in HCT116-shLuc and HCT116-shPG cells cultured in regular medium (-CM), conditioned medium from unstimulated fibroblasts (CM N-AF), or conditioned medium from progastrin-stimulated fibroblasts (CM PG-AF).

Because the response of tumour cells to fibroblast-derived chemokines may also depend on receptor expression, we analysed CXCR4, CXCR1 and CXCR2 mRNA levels in HCT116-shLuc and HCT116-shPG cells exposed to conditioned medium from unstimulated fibroblasts (CM N-AF) or progastrin-stimulated fibroblasts (CM PG-AF). Basal receptor expression did not differ between HCT116-shLuc and HCT116-shPG cells. In HCT116-shLuc cells, fibroblast-conditioned media increased CXCR4 expression relative to regular medium, but no significant difference was observed between CM N-AF and CM PG-AF. In HCT116-shPG cells, CM PG-AF tended to increase CXCR4 expression without reaching significance. CXCR1 expression was induced by fibroblast-conditioned media in both cell populations and was significantly higher in HCT116-shPG cells exposed to CM PG-AF than to CM N-AF, whereas CXCR2 regulation was limited (Fig. 4b-d). These data support a model in which progastrin-activated fibroblasts enhance tumour cell migration mainly through paracrine CXCL12 and CXCL8 signalling, potentially accompanied by modulation of chemokine receptor expression in epithelial cells.

### Progastrin activates PI3K/Akt signalling in fibroblasts and PI3K inhibition suppresses chemokine secretion and fibroblast-dependent migration

We previously reported that progastrin activates PI3K/Akt signalling in colonic epithelial cells [29], and Duckworth et al. implicated the same pathway in progastrin-induced fibroblast proliferation [22]. We therefore examined whether PI3K/Akt signalling mediates the pro-migratory secretory response observed here. In CCD18Co fibroblasts, progastrin induced sustained Akt phosphorylation detectable from 3 min to 4 h after stimulation, whereas total Akt levels remained unchanged (Fig. 5a).

**Figure 5.**
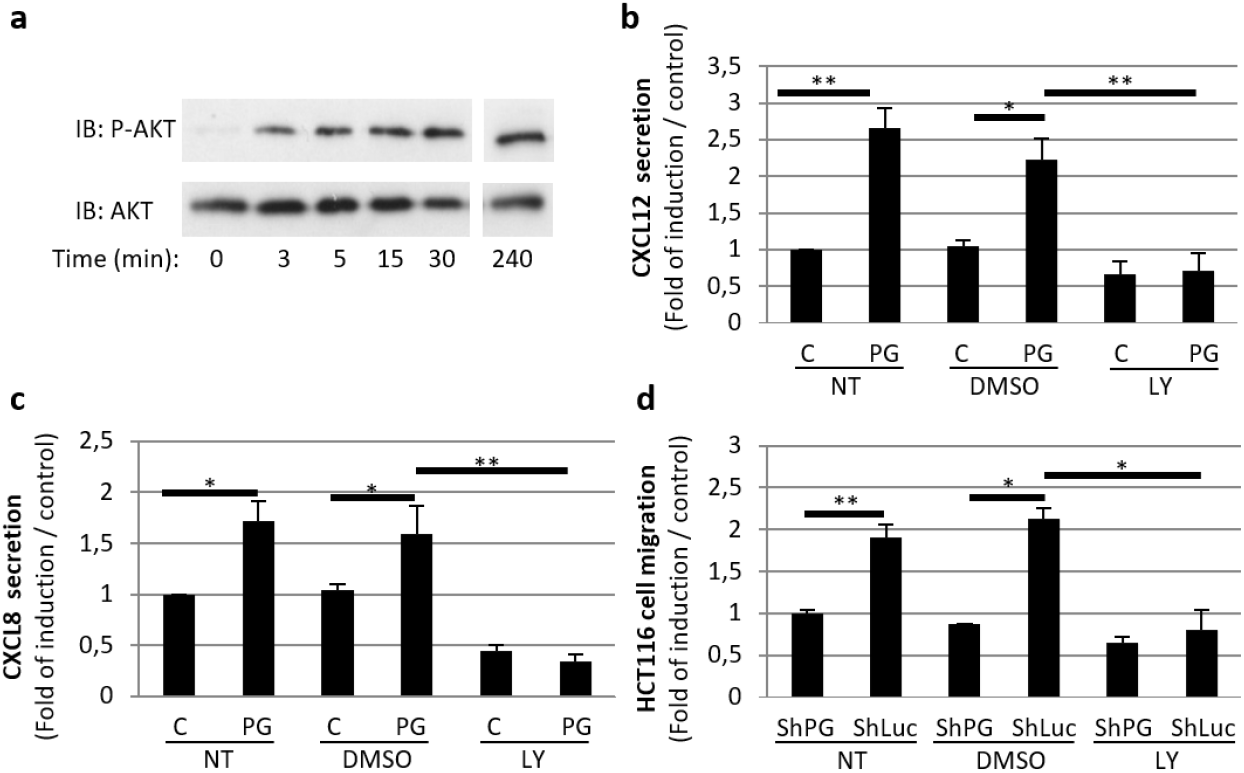
PI3K/Akt signalling is required for the chemokine secretory response and pro-migratory activity of progastrin-stimulated fibroblasts. **a**, Western blot analysis of phospho-Akt (Ser473) and total Akt in CCD18Co fibroblasts stimulated with progastrin for the indicated times. **b-c**, ELISA quantification of CXCL12 and CXCL8 secretion in CCD18Co-conditioned medium after treatment with progastrin in the presence of DMSO or LY294002. **d**, Transwell migration assays showing the effect of fibroblast PI3K inhibition on HCT116 cell migration.

To assess the functional importance of this pathway, CCD18Co cells were pretreated with LY294002 before progastrin exposure. PI3K inhibition markedly reduced progastrin-induced CXCL12 and CXCL8 secretion, as measured by ELISA (Fig. 5b-c). Importantly, when fibroblasts were pretreated with LY294002 before coculture with HCT116 cells, the fibroblast-dependent migratory effect was completely suppressed (Fig. 5d). These results indicate that PI3K/Akt signalling in fibroblasts is required for the chemokine secretory programme elicited by progastrin and for the resulting enhancement of CRC cell migration.

## Discussion

The present study identifies a stromal mechanism by which progastrin can potentiate CRC progression. Progastrin has long been viewed primarily as a trophic factor for the intestinal epithelium and for CRC cells [14–19]. Here, we show that it also acts on resident colonic fibroblasts, inducing a CAF-like activation state characterized by increased FAPα and αSMA expression and the acquisition of a secretory programme that enhances tumour cell migration. The work therefore extends the biological scope of progastrin from epithelial growth control to active remodelling of the tumour microenvironment.

The evidence that progastrin activates fibroblasts is supported by both in vivo and in vitro observations. In hGAS mice, stromal FAPα and αSMA expression were readily detectable in the colonic mucosa, whereas control tissue lacked these activation markers. In CCD18Co cells, progastrin directly induced both proteins. These data are consistent with the concept that tumour-derived factors can convert resident fibroblasts into myofibroblastic, tumour-supportive stromal cells [25,26]. They also complement the study by Duckworth et al., who showed that progastrin stimulates colonic myofibroblasts and promotes epithelial proliferation through IGF-II secretion [22]. Our data indicate that the stromal consequences of progastrin extend beyond proliferative support to include induction of a pro-migratory paracrine programme.

The functional core of the study is the demonstration that tumour cell migration becomes strongly dependent on fibroblasts and is further amplified when tumour cells express progastrin. This effect could be transferred by preconditioning fibroblasts with progastrin-producing tumour cells, arguing for a genuine stromal memory of exposure. The experimental endpoint measured here is migration, not invasion through extracellular matrix or metastatic colonization. The data therefore support a fibroblast-dependent pro-migratory mechanism rather than direct proof of metastasis. Nonetheless, this phenotype is highly relevant biologically because fibroblast activation is a recognized prerequisite for invasive conversion in CRC [5–7,26].

A mechanistic strength of the work is the identification of CXCL12 and CXCL8 as key mediators of this stromal effect. Both chemokines are well implicated in CRC biology. CXCL12 signalling through CXCR4 promotes tumour cell dissemination, while CXCL8 contributes to tumour progression, angiogenesis and metastatic competence [27,28]. We show that progastrin induces expression and secretion of both chemokines by fibroblasts, and that neutralization of either one suppresses the additional migratory advantage conferred by progastrin-activated fibroblasts. The chemokine receptor analysis further suggests that fibroblast-derived signals may sensitize epithelial cells to this paracrine environment, particularly through CXCR1 induction in low-progastrin cells. Taken together, the data support a feed-forward model in which tumour-derived progastrin transforms fibroblasts into chemokine-secreting stromal partners that reinforce tumour cell motility.

Our results also position PI3K/Akt as a central intracellular relay in fibroblasts downstream of progastrin. This pathway had already been linked to progastrin signalling in epithelial cells [29] and to fibroblast proliferation [22]. Here, progastrin induced sustained Akt phosphorylation in CCD18Co cells, and pharmacological PI3K inhibition suppressed both CXCL12/CXCL8 secretion and fibroblast-dependent tumour cell migration. These findings provide a mechanistic framework for the stromal response to progastrin and suggest that the peptide engages signalling pathways shared by both epithelial and stromal compartments, albeit with distinct phenotypic outputs.

We acknowledge that the mechanistic work relies on a single normal fibroblast line and a single CRC cell line. However, despite this limitation, the overall coherence of the *in vivo*, biochemical, transcriptional and functional data supports the main conclusion that progastrin is not only an epithelial trophic factor, but also a stromal reprogramming signal in CRC.

In conclusion, this work identifies progastrin as a mediator of tumour-stroma crosstalk that activates colonic fibroblasts and elicits a PI3K/Akt-dependent CXCL12/CXCL8 secretory programme promoting CRC cell migration. These findings provide a mechanistic explanation for how a peptide classically associated with epithelial proliferation may also contribute to tumour cell motility through the stromal compartment. They further reinforce the translational interest of targeting extracellular progastrin in CRC, not only to restrain tumour cell-autonomous growth signals, but also to disrupt a pro-migratory stromal niche.

## Acknowledgments

This work was funded by the Plan Cancer project Mocassin-Biosystem 2017 granted to AF and supported by the INSERM.

---

**Supplementary Figure S1.**
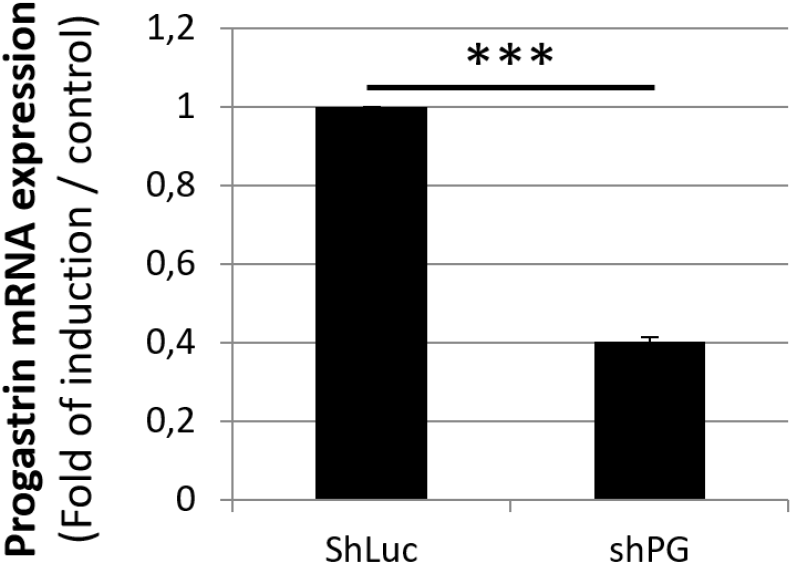
Validation of progastrin knockdown in HCT116 cells. RT-qPCR analysis of GAST/progastrin mRNA expression in HCT116-shLuc and HCT116-shPG cells.

